# Data-Poor Ecological Risk Assessment of Multiple Stressors

**DOI:** 10.1101/2021.11.11.468297

**Authors:** Richard E Grewelle, Elizabeth Mansfield, Fiorenza Micheli, Giulio De Leo

## Abstract

1. Ecological Risk Assessment is a formal process widely applied to terrestrial, marine, and freshwater ecosystems to evaluate the likelihood of adverse ecological effects occurring as a result of exposure to natural or anthropogenic stressors. For many species, data is sparse and semi-quantitative methodologies provide valuable insight for ecosystem management. Recent statistical developments have improved the quality of these analyses yet a rigorous theoretical framework to assess the cumulative impact of multiple stressors is lacking.
2. We present EcoRAMS, a web application and open-source software module that provides easy-to-use, statistically-robust ecological risk assessments of multiple stressors in data-poor contexts. The software receives attribute scores for two variables (e.g. exposure-sensitivity, productivity-susceptibility, severity-likelihood) via CSV templates and outputs results according to a probabilistic metric of risk.
3. We demonstrate comparative results across a range of assumptions, using simulated and empirical datasets including up to five stressors. Accounting for multiple stressors even when data is limited provides a more detailed analysis of risk that may otherwise be understated in single stressor analyses.
4. This application will allow quantification of risk across data-poor contexts for which statistical results have been previously unavailable. The web app format of EcoRAMS.net lowers the barrier of use for practitioners and scientists at any level of statistical training.

## Introduction

A wide variety of approaches are deployed to measure risk of species to stressors, which are often human-mediated. These stressors can be unique to each ecosystem or even to species within an ecosystem. Common stressors of wildlife result from harvesting, mining, recreational, or construction activities and those chosen for analysis are relevant to the species affected and stakeholder interests (Hope, 2006). In some contexts, the effects of stressors is quantified because the response of a species or endpoint can be measured as the strength of the stressor is titrated (e.g. increasing concentration of a contaminant) (Norton et al., 1992). However, in many contexts, the relationship between stressor and response is unknown or under-studied. Semi-quantitative methodologies have been developed to assess risk with incomplete data and often rely on expert input and standardized scoring procedures (Pilling et al., 2009). This is particularly common in the marine environment for fished and bycatch species because data collection is labor and resource intensive. Frameworks like the Productivity-Susceptibility Analysis (PSA) have been incorporated within the broader Ecological Risk Assessment for the Effects of Fishing (ERAEF) approach which uses risk estimates from the PSA for downstream assessment by managers, scientists, and stakeholders (Hobday et al., 2011; Hobday et al., 2007; Stobutzki et al., 2001). Management priorities can account for all species, guilds, and communities with these approaches and is the goal of ecosystem based management (Hazen et al., 2016; Townsend et al., 2019). For this reason, data-poor methodologies have been widely adopted (Battista et al., 2017; Marine Stewardship Council, 2019; Pontón-Cevallos et al., 2020). However, data-poor ecological risk assessments, like the PSA and others employing Exposure and Sensitivity or Effect as risk determinants, have historically lacked a statistical basis.

In our previous work, we identified several sources of bias in the widely used PSA due primarily to the changing distribution of species that occurs under different analysis conditions, such as model choice and number of attributes used. Because a theoretical basis for the PSA was absent, we provided a comprehensive probabilistic interpretation that eliminates bias in all forms of the analysis (Grewelle et al., 2021). The new rPSA framework is able to incorporate additive and multiplicative models of Productivity or Susceptibility and addresses changes in the distribution of species’ scores on the two-dimensional PSA plot as the number of attributes used increases. We showed that both an untransformed additive model and a log-transformed multiplicative model produce a normal distribution. We then provided an analytical formulation of Vulnerability by projecting the resulting bivariate normal distribution onto a one-dimensional risk axis (Fig. 1).

**Figure 1:**
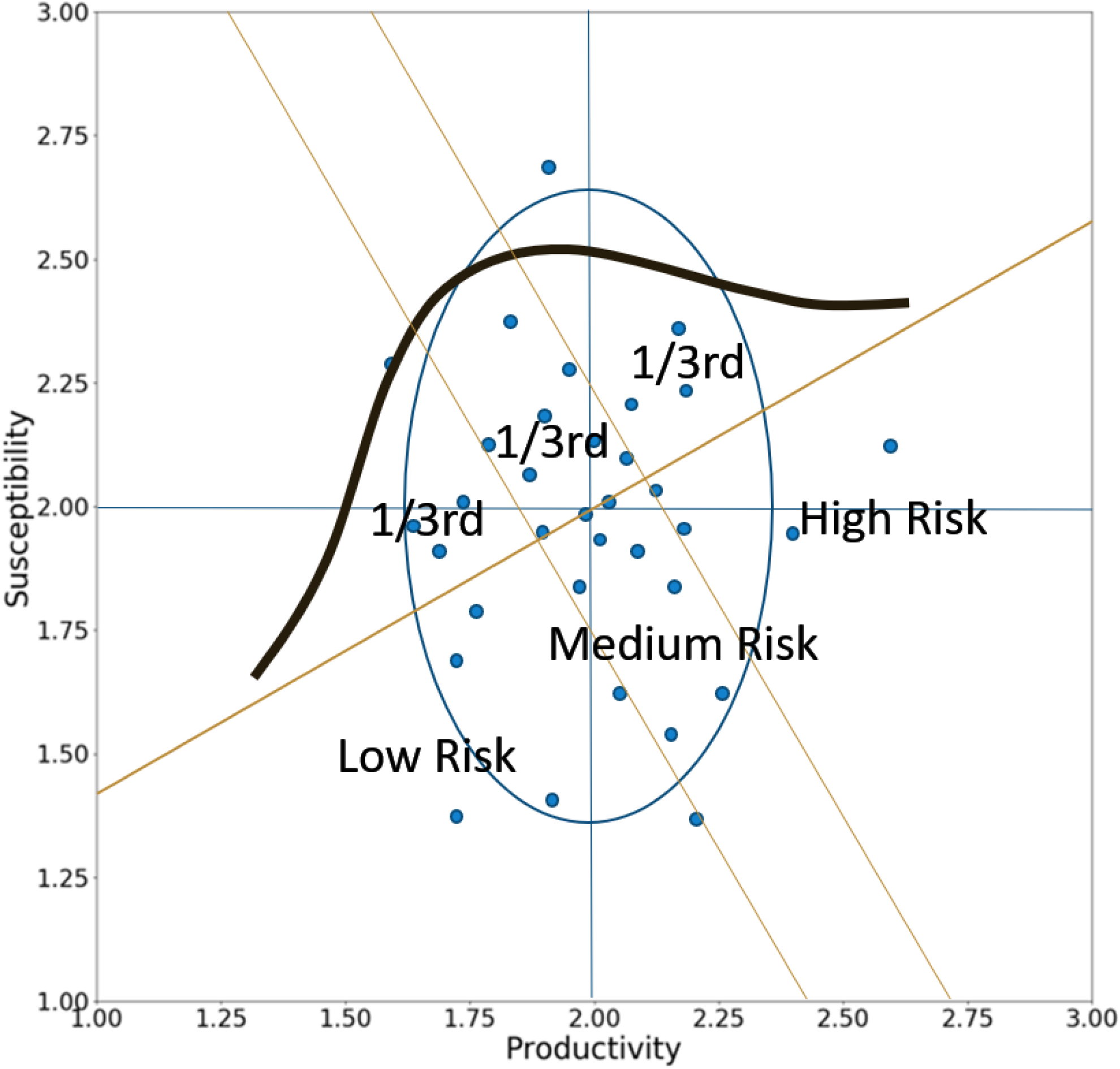
In the scoring model used by the PSA, a bivariate normal distribution (blue ellipse) is produced along the Productivity and Susceptibility axes. This can be projected along a Risk Axis (bold orange, ascending left to right), which is defined by the likelihood properties of the analysis, to form an ordered set of points along a one dimensional normal distribution (bold black). This new distribution is used to score Vulnerability. Vulnerability scores fall into risk categories delineated by thresholds (light orange, descending left to right) which divide the distribution of points into equal partitions by probability.

Vulnerability in our previous work is presented in two forms: a probability (*V*_*p*_) taking values between 0 (lowest risk) and 1 (highest risk), which are comparable between studies, and a relative value (*V*) that is easily calculated and is useful to categorize and rank species within a study but cannot be compared between studies with different assumptions. These formulations of Vulnerability rely on precise characterization of the distribution of expected scores on both axes of the PSA plot, allowing both *V*_*p*_ and *V* to be statistical quantities.

The PSA as designed for data-poor, small scale fisheries assesses Susceptibility to a single ecological stressor, i.e. a specific fishery or fishing gear. However, in many ecological risk assessments, species are subject to multiple stressors (Halpern et al., 2009; Van den Brink et al., 2016). Building on the original PSA framework, Micheli et al. presented an empirical formulate to compute an Aggregated Susceptibility (AS) that broadened the definition of Susceptibility to multiple fishing gears or stressors (Micheli et al., 2014). Aggregated Susceptibility is truncated at a maximum value of 3 to remain within the scoring bounds of the PSA.

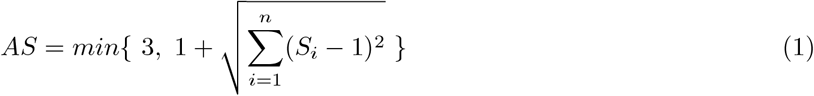

Subtracting 1 from each Susceptibility score standardizes the procedure of squaring, summation, and taking the square root of the internal expression. When Susceptibility to a stressor is 1, AS does not increase. Adding 1 to this expression allows AS to take a minimum value of 1. Therefore, the AS adheres most closely to a variant calculation for Vulnerability

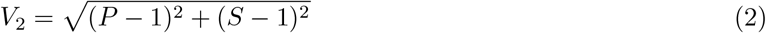

rather than the earlier expression:

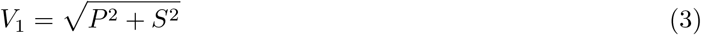

Each of these variants is heuristical and neither possesses a theoretical basis nor is unbiased, hence the need to provide a basis with the rPSA. Though practical, the empirical formula to calculate AS was not derived on the basis of rigorous statistical principles. The primary aim of this work is to create a statistical interpretation of AS that can be integrated with the rPSA framework which will produce a robust measure of Vulnerability for when one or more stressors contribute to species’ Susceptibility. We generalize this framework for data-poor ecological risk assessment to evaluate any two-dimensional scoring procedure, including Sensitivity-Exposure or Severity-Likelihood based analysis.

## Methods

All variables scored by calculating the mean of several attributes are normally distributed, and a full treatment of these statistics is available (Grewelle et al., 2021). Calculation of Vulnerability with the rPSA requires that each variable is normally distributed, and the mean and standard error must be known. We denote *α* as any response variable (Productivity, Sensitivity, Effect, Severity, etc.) determined by characteristics of the species or endpoints studied and *β* as any stressor variable (Susceptibility, Exposure, Likelihood, etc.) determined by the probability of contact with stressors. It is assumed that *β* is a composite variable with basis {*β*_1_, *β*_2_, …, *β*_*n*_}, while *α* is not. Though, this method can be generalized to relax this assumption. We modify the equation for AS to a statistical form to calculate *β. β* must be Gaussian; however, its current form is not a statistical distribution. Each stressor variable composing *β* is Gaussian because they are scored by calculating the mean of several attributes. With modification we can standardize the expression within the square root of AS to form a chi distribution:

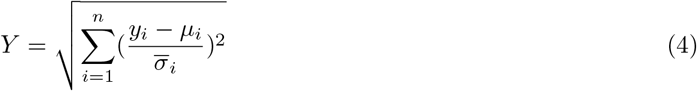

There are *k* species or endpoints evaluated with *n* stressor variables. *y*_*i*_ is the score of each stressor variable distributed according to 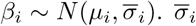. is the standard error of the mean expected score determined by scoring criteria. *Y* is a statistic that is chi distributed

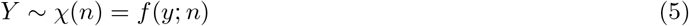

with cumulative distribution function *F*_+_(*y*; *n*). This function gives *Pr*(*Y* ≤ *y*) where *Y >* 0. Each term in the summation of Y is a squared Z-score, and the sign of each Z-score is not preserved. We form a piecewise function *F* which translates the sign and magnitude into a continuous cumulative distribution function on *y* ∈ (−∞, ∞). Let 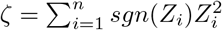, where *Z*_*i*_ is the Z-score of each stressor variable.

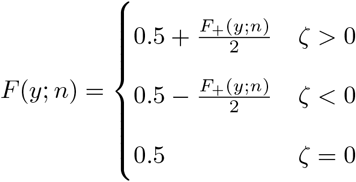

This result follows from the additive property of the chi distribution:

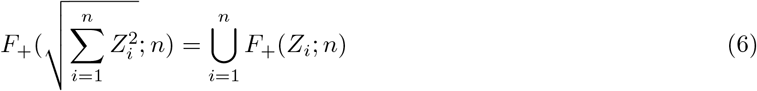

*f* (*y*; *n*) is a symmetric probability density function centered at *y* = 0. To integrate this distribution with the rPSA, we transform the modified chi distribution to a normal distribution centered at *y* = 2 with a chosen variance. By the probability integral transform theorem, we can map the chi-distributed points {*p*} to a normal distribution with mapping *m*({*p*}):

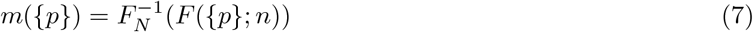

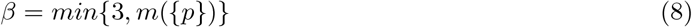

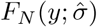 is a normal cumulative distribution function whose standard error, 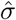, can be chosen arbitrarily because the mapped points and this standard error are used to generate Vulnerability statistics when incorporated in the rPSA framework, which standardizes values by any chosen standard error. For visualization we choose to standardize the calculation of 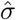 to be similar in magnitude to the standard errors of each contributing stressor variable. For those variables log-transformed to produce a normal distribution, corresponding standard errors can be adjusted for subsequent calculation using the lognormal relation:

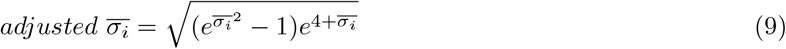

Additive stressor variables require no transformation, and therefore their associated adjusted standard errors are equivalent to their standard errors without adjustment. We propagate error for the mean of the stressor variables, assuming variables are independent of one another:

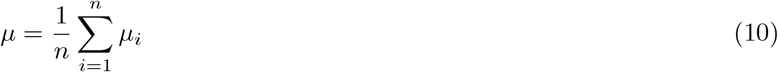

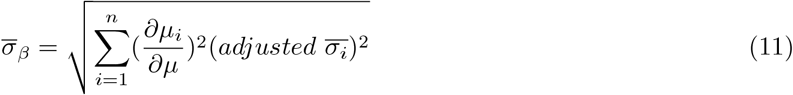

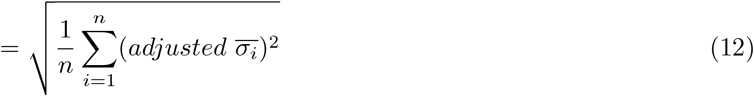

The probability density function for the normal mapping is 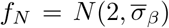. Recalling from (Grewelle et al., 2021) any vector 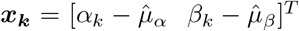 can be projected along the risk vector, ***r***, to map ***X*** to one dimension. *k* is the species index. After mapping all points to the risk vector, the distance of each point *x*_*k*_ = (*α*_*k*_, *β*_*k*_) from the mean is the magnitude of the projection:

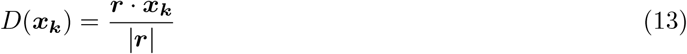

The projection results in a linear transformation of ***X*** with standard error

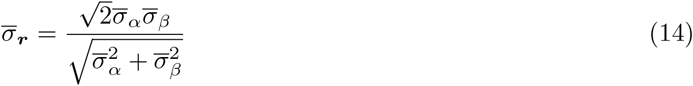

The cumulative distribution function that yields the probabilistic metric of Vulnerability is:

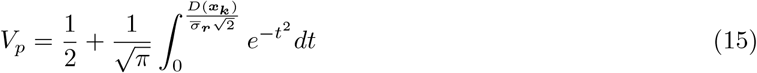

This metric can be compared across all varieties of input conditions, including variable number and type of stressors and attributes. However, comparisons across studies are most explicable when conditions of the analyses are similar.

### Compounding Stressors

The above framework implemented in EcoRAMS ([data-poor] Ecological Risk Assessment of Multiple Stressors) measures Vulnerability from multiple stressor variables, standardized by the distance of each stressor score from each mean set by the attribute scoring procedure and the chosen model (additive or multiplicative). Vulnerability values calculated in this way will fall between Vulnerability values calculated for each stressor independently, and therefore, this metric provides maximum resolution to distribute points across the low, medium, high risk range. This interpretation of Vulnerability assumes that stressors hold values close to their mean when not included in analysis (i.e. the null model is the mean for each stressor). Nevertheless, it may be useful to assume that stressors hold values close to their minimum when otherwise not included in analysis (i.e. the null model is the minimum for each stressor). In these instances, Vulnerability should increase as stressors are added to the analysis. In fact, this is the impetus for the concept of Aggregated Susceptibility. EcoRAMS incorporates compounding stressors by calculating the standardized distance of each score from a minimum to form the inner expression of *Y*.

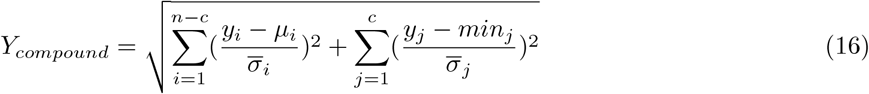

*c* is the number of stressor variables to be compounded. 1 ≤ *n* − *c* because at least one variable must be standardized at the mean, not the minimum. For an additive model, the default minimum is set to 1 as attribute scores take values between 1 and 3. In the multiplicative model, the default minimum is *log*(1) = 0.

### Scoring Practices

The EcoRAMS method requires several steps prior to submitting template CSV files to receive results (Fig. 2). After the analysis conditions are setup, the *α* and *β* attributes must be chosen. Attributes should be approximately independent from each other. Redundant attributes should be removed, or otherwise, the effective number of attributes can be estimated using the method outlined in the supplementary information of (Grewelle et al., 2021). Percentiles are then chosen for scoring each attribute, adhering to relevant expected ranges. For ease, we recommend equal bin sizes: 1 = 0-33%, 2 = 33-67%, 3 = 67-100%. When attributes are irrelevant or lack expert input or data for scoring, the attributes should not be scored. Leaving the corresponding cells blank in the CSV templates provided will adjust the number of attributes used for the species and broaden standard error estimates for calculating Vulnerability and scored attributes will be appropriately weighted. The *α* and *β* values are calculated as the weighted means of their respective attribute scores. Weights can be applied on the basis of importance or data quality, and efforts to justify weights via supporting mechanism or data should be made. The mean calculated depends on the model assumed for the attributes (geometric mean = multiplicative, arithmetic mean = additive). However, as a default, *α* variables should be additive, while *β* variables should be multiplicative. A multiplicative model should be used when the magnitude of risk associated with a variable depends on interactions of the attributes with each other such that absence of risk for any attribute substantially reduces risk as a whole. Values of variables scored in this way act much like probabilities, and in fact, a geometric mean is an isotone (order-preserving) mapping of probability and is indistinguishable in the results of the EcoRAMS analysis. The model used can be different for each variable *β*_1_, *β*_2_, …, *β*_*n*_ if different attributes are used to score each, though we encourage consistency for ease of interpretation. Not all risk analysis are amenable to the above scoring procedure, and if not, EcoRAMS should not be used. Risk analyses scored differently may be adapted to use with EcoRAMS by defining scoring criteria within the range of 1-3.

**Figure 2:**
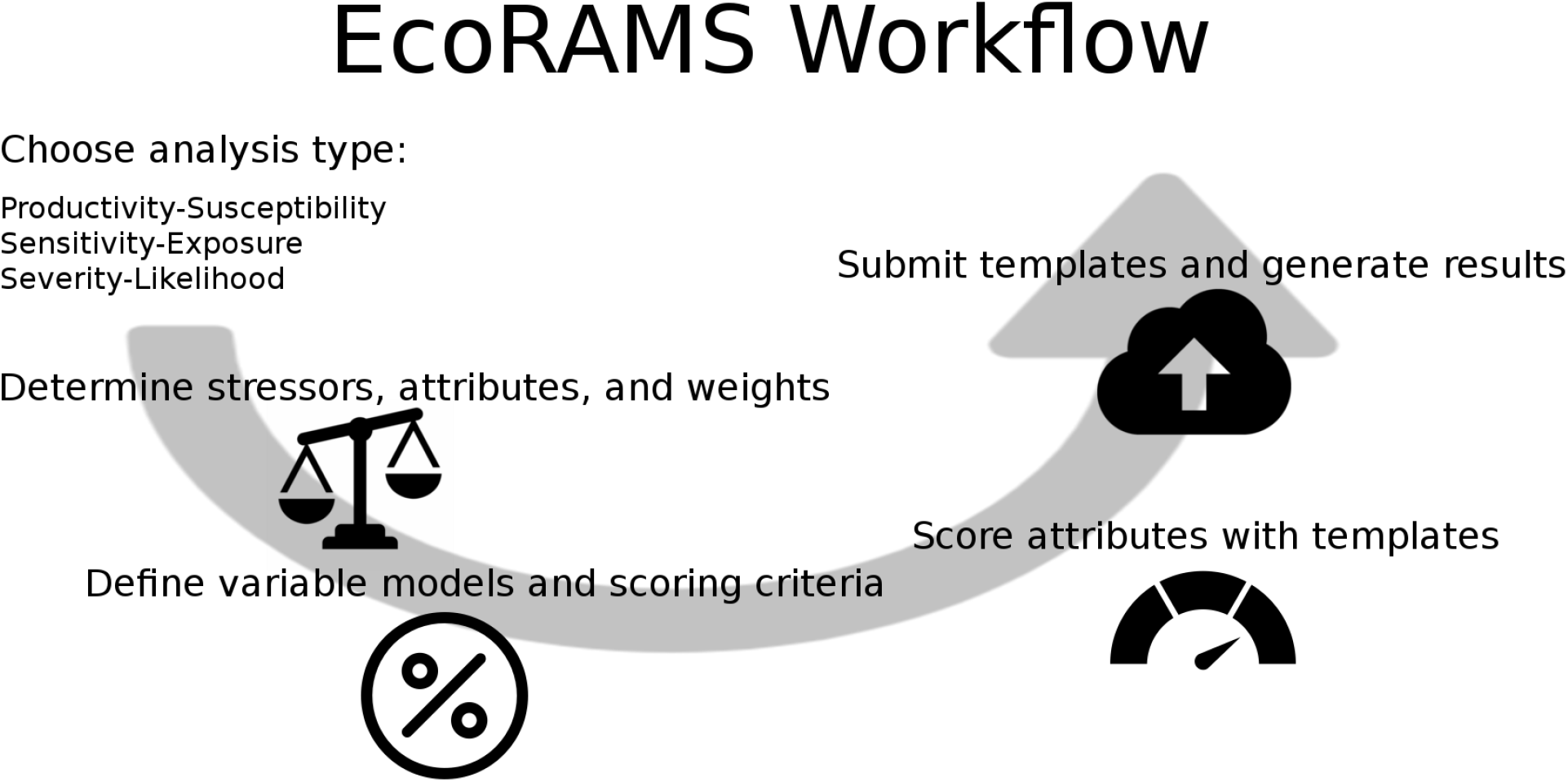
Workflow diagram for EcoRAMS analysis. The analysis type chosen determines which of the pairs of variables is used in the risk assessment, which will constrain the types of attributes used for scoring. Included stressors, attribute weights, and their associated models and scoring criteria will define the scope and structure of the analysis. After attributes are scored using provided templates, users submit templates simultaneously on the EcoRAMS web app at EcoRAMS.net and results are automatically generated.

### Case Studies

To demonstrate the use of EcoRAMS, we generated a set of data for 100 species using a standard scoring procedure for the PSA used in the Ecological Risk Assessment for the Effects of Fishing. We adopt the same scoring procedure by which attributes of *α* and *β* variables are scored 1 (low risk), 2 (medium risk), or 3 (high risk). Note that these scores reflect association with risk, not association with the *α* or *β* variables per se. For example, in a PSA high Productivity is associated with low risk (Hobday et al., 2007), and therefore Productivity attribute scores should be transformed by subtracting from 4 if originally scored as 1 = low Productivity, 2 = medium Productivity, 3 = high Productivity.

In our simulated analysis, 7 *α* and 4 *β* attributes are scored randomly on a uniform distribution of integers between 1 and 3. The *α* score is calculated as the arithmetic mean assuming an additive model. The *β* score is calculated as the geometric mean assuming a multiplicative model. Attributes are weighted equally. Fig. 3 presents the results of EcoRAMS when *β* variables are not compounded for 1-3 stressors, while Fig. 4 presents results when stressors 2 and 3 compound on stressor 1.

**Figure 3:**
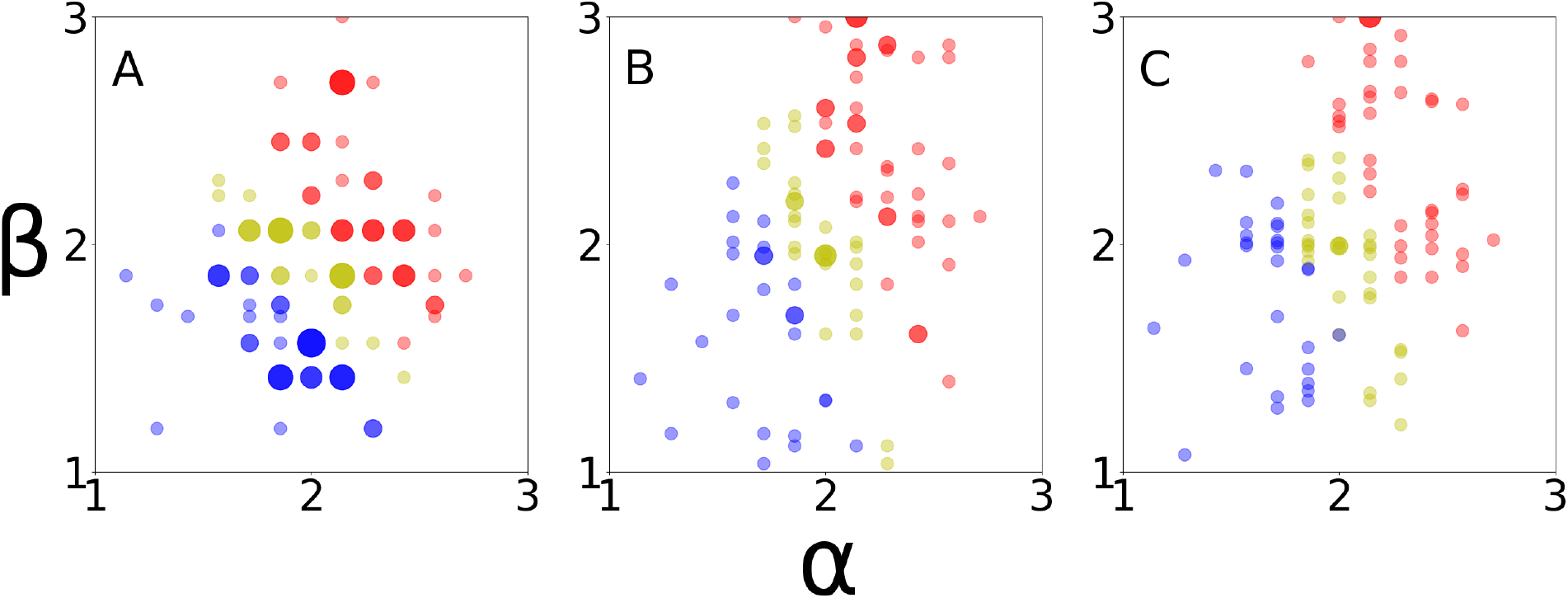
Risk classification of 100 simulated species by Vulnerability scores in which stressors do not compound. Low risk species are in blue, medium risk in yellow, high risk in red. The size of a dot corresponds to the number of species sharing overlapping positions in the plot. (A) A single stressor (*β*_1_) is used in an analysis. (B) Two stressors are used in an analysis, the second (*β*_2_) does not compound on the first shown in panel A. (C) Three stressors (*β*_1_, *β*_2_, *β*_3_) are used in an analysis.

**Figure 4:**
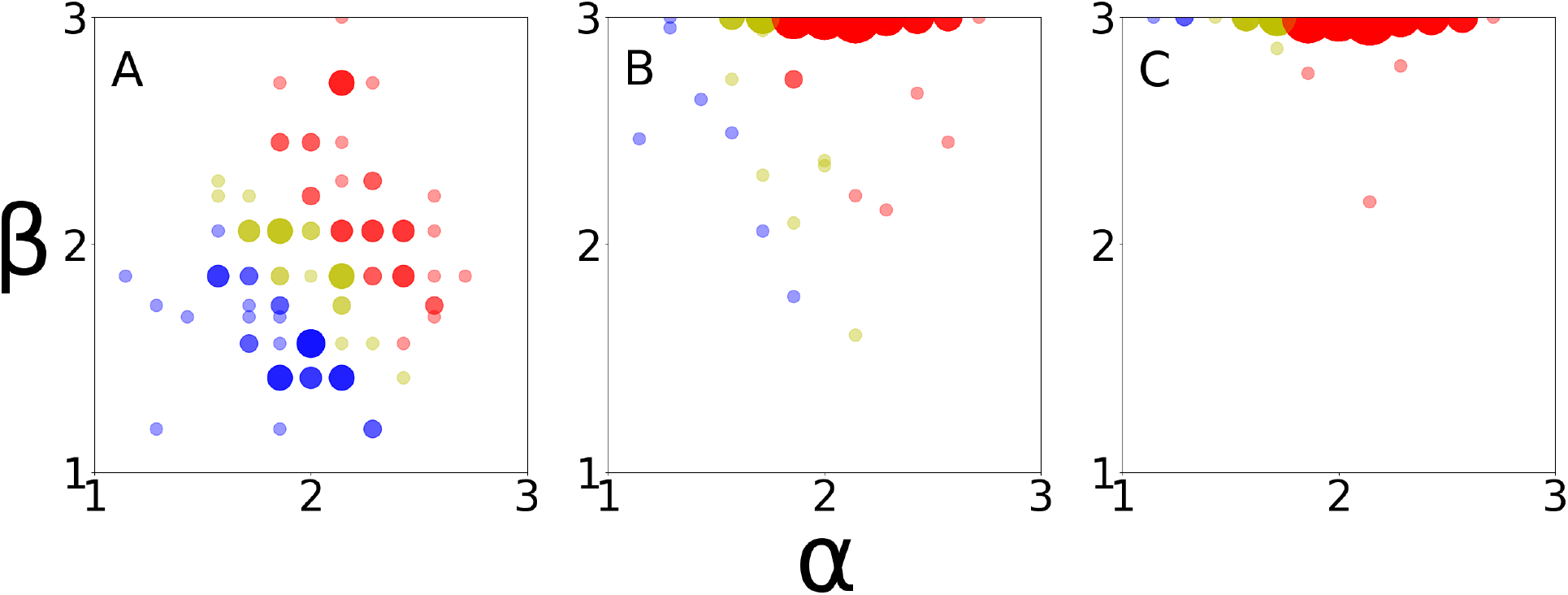
Risk classification of 100 simulated species by Vulnerability scores in which stressors compound. Low risk species are in blue, medium risk in yellow, high risk in red. The size of a dot corresponds to the number of species sharing overlapping positions in the plot. (A) A single stressor (*β* variable) is used in an analysis. (B) Two stressors are used in an analysis, the second (*β*_2_) compounds on the first shown in panel A. (C) Three stressors (*β*_1_, *β*_2_, *β*_3_) are used in an analysis, the second and third compound on the first.

When stressors do not compound and are instead standardized by the expected mean, additional stressors will not affect the distribution of risk categorization. When scoring is uniformly distributed, approximately one-third of species should fall into each category. This procedure is useful for maximally discriminating risk of species within a study for downstream assessment. However, in some studies when it assumed that stressors not directly measured are negligible, the addition of stressor variables presents an opportunity to assess cumulative impacts of two or more stressors. In these cases, Vulnerability scores are expected to increase with added stressors for which scores exceed the minimum, and risk compounds. Fig. 4 shows how the distribution of risk rapidly shifts as stressors compound and Vulnerability scores increase.

When a minimum score is used to standardize additional stressors, fewer species are expected to remain in the low or medium risk categories, especially with three or more stressors. *β* scores are truncated at 3, which ensures that some species will remain in these categories when *α* scores are sufficiently low. In many scenarios a far greater number of species will remain in lower categories than depicted in Fig. 4 because scoring of additional stressors will be skewed right, favoring lower scores than the stressor of greatest impact, which may be the initial target of analysis. We see this illustrated in an empirical case study of 81 fished species in Baja California, Mexico. We re-analyzed these species from (Micheli et al., 2014) using the EcoRAMS framework (Fig. 5). Each type of fishing activity (e.g. gear type) is treated as a separate stressor.

**Figure 5:**
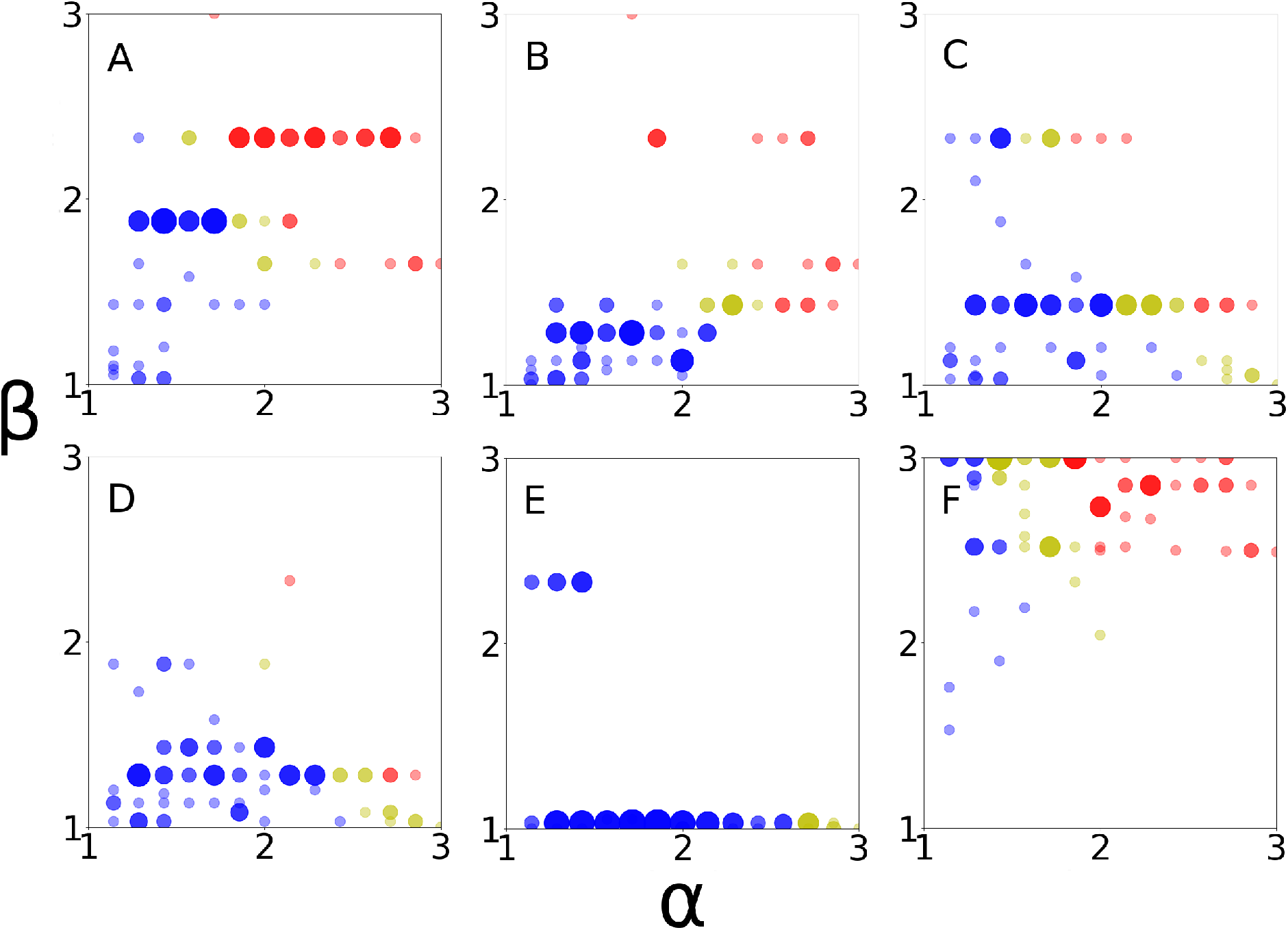
Risk classification of 81 marine species in Baja California, Mexico analyzed by Micheli et al. Low risk species are in blue, medium risk in yellow, high risk in red. The size of a dot corresponds to the number of species sharing overlapping positions in the plot. *α* = Productivity and *β* = Susceptibility. (A) Analysis of stressor of highest impact, set gillnets, followed by analyses of four other fishing stressors in descending order of impact: (B) drift gillnets, (C) lobster traps, (D) fish traps, (E) dive fishing. (F) These stressors are analyzed together in the EcoRAMS framework assuming stressors compound, revealing higher Vulnerability scores due to increased Susceptibility than for any stressor alone.

Despite the majority of species receiving Vulnerability scores placing them in the low risk category for each single stressor (Fig. 5 A-E), the cumulative impacts of each of the five stressors produces a higher Susceptibility score, and therefore, a higher Vulnerability score when compounded. Compared with the single stressor of highest impact (set gillnets), accounting for multiple stressors results in 26% higher net risk: 23% low, 31% medium, 46% high for multiple stressors compared to 49% low, 10% medium, 41% high for set gillnets. Although this analysis highlights application to multiple sources of fishing pressure, EcoRAMS is fully flexible to analyze any system for which multiple independent sources of stress introduce risk to subjects of the analysis. For example, the PSA has been used to evaluate terrestrial ecosystems, although it was designed for data-poor fisheries. Similar risk-based frameworks that use Exposure and Sensitivity (or Effect) or Severity and Likelihood as variables have been applied to marine and terrestrial systems and have also been extensively developed for data-rich ecological risk assessments (Council et al., 2009; Samhouri et al., 2019). These applications extend beyond ecology and include analysis of risk in business (Koller, 2005), human health (Ravindra & Mor, 2019; World Health Organization et al., 2020), engineering (Zio, 2018), and others. We provide a cohesive statistical framework, associated software, and web application to easily and robustly analyze risk in any data-poor context, including when accounting for multiple stressors is important for the assessment. EcoRAMS requires no prior statistical or programming knowledge, as full functionality of this software is deployed as a web app that only requires users to complete template CSV files with their data. We view wide and reliable access to robust statistical methods as imperative to the progress of ecosystem-based management and risk analysis. Because 90+% of marine species are considered data-poor (FAO, 2020; Mora et al., 2011), EcoRAMS has the opportunity to be widely adopted to improve analyses marine sciences, and we anticipate EcoRAMS to have similar value to other ecological risk assessments of terrestrial and freshwater systems when data is sparse. The best use of EcoRAMS is with input from scientists, managers, practitioners, and stakeholders to allocate time and resources to species for which conservation provides mutual benefits to the ecosystem and the people within it (Finkbeiner et al., 2017; Oestreich et al., 2019).

### Considerations

Statistical (probabilistic) methods have been deployed for ecological risk assessments where data is available. We provide a statistical framework to conduct ecological risk assessments in data-poor contexts. Even in data-rich contexts, the use of statistical risk assessments is not universally accepted practice. Challenges like increased complexity, greater data needs, and difficulty in communicating results to stakeholders slow widespread uptake (Hope, 2006). Conventional non-statistical methods like hazard quotients or guidelines set to fixed thresholds without regard to uncertainty in data or mechanistic interpretation of the underlying model are often simple and convenient interpretations of risk and are used by decision-makers (Tannenbaum et al., 2003), but these benefits become less tenable when statistical practices are highly accessible. Changing convention will take time, and ecological risk assessments will benefit from the transition to statistical methods, as they provide more reliable insights into risk management. Adoption of statistical methods for data-poor ecological risk assessments may similarly require time, and they key to improving the state of the field in the shortest time is to make methods accessible. EcoRAMS represents a sophisticated statistical software with easy inputs and easily understood results for any audience. Engagement with stakeholders and downstream efforts to prioritize management is crucial for any risk assessment, and EcoRAMS can facilitate these synergies.

## Acknowledgements

We thank all members of the De Leo lab for support on this project. REG is funded by the Stanford Graduate Fellowship and the ARCS Fellowship. EJM is funded by the Joseph R. McMicking Fellowship for Biological Sciences. The project was supported by NSF grant #1736830.

## Authors’ Contributions

Conceptualization, REG; Methodology, REG; Software, REG; Analysis, REG & EM; Visualization, REG; Writing – Original Draft, REG; Writing – Review & Editing, REG, EM, FM, GDL; Funding Acquisition, REG, FM, GDL

## Data Availability

Software associated with this study is available for download and use at https://github.com/grewelle/DPERAMS. Empirical data evaluated is publicly available in referenced works. The web app is deployed at EcoRAMS.net.

## Supplement

Downloadable templates and instructions for EcoRAMS, also found at https://github.com/grewelle/EcoRAMS/blob/main/README.md

EcoRAMS is software used to perform ecological risk assessments in data-poor contexts. It provides as statistical interpretation of risk for each species or endpoint analyzed. This metric is probabilistic Vulnerability (Vp). Inputs are attribute scores and weights for two variables: response (e.g. Productivity, Effect, Sensitivity, Severity) and stressor (e.g. Susceptibility, Exposure, Likelihood). EcoRAMS is designed to incorporate multiple stressors that when aggregated can have compound impacts on risk or not. This software is associated with Grewelle et al. 2021, Data-Poor Ecological Risk Assessment of Multiple Stressors. This paper builds on previous work which constructs a statistical framework for data-poor risk assessments in fisheries with single stressors, Grewelle et al. 2021, Redefining Risk in Data-Poor Fisheries https://onlinelibrary.wiley.com/doi/abs/10.1111/faf.12561, https://github.com/grewelle/rPSA and earlier work on multiple fishery stressors Micheli et al. 2014, A risk-based framework for assessing the cumulative impact of multiple fisheries https://web.stanford.edu/group/MicheliLab/pdf/a%20risk%20based%20framework.pdf. EcoRAMS is free to use, and we welcome feedback to improve the usability of the software. We only ask that you credit our work and cite our papers in any byproducts resulting from use of EcoRAMS.

Below is a guide to using EcoRAMS. After downloading and completing templates, analysis occurs within a few seconds of upload. If your data is well organized to be input into templates, a full analysis from template download to results can occur within a few minutes. The instructions are divided into three sections: pre-download of templates, template completion, and EcoRAMS analysis.

### 1. Pre-download of templates

- Determine the types of response and stressor variables used in analysis (e.g. Productivity-Susceptibility, Exposure-Effect/Sensitivity, Severity-Likelihood) - Generate a set of one or more stressor variables that independently contribute to risk - Decide which set of attributes will be used to assess each of the variables. If more than one stressor is included, different attributes may be used for different types of stressors, though care should be taken to interpret results appropriately given the added complexity of the analysis. Chosen attributes should be approximately independent from each other. Highly correlated/redundant attributes should not be included in the analysis. If moderately-highly correlated attributes are used, refer to the supplement of Grewelle et al. 2021, *Redefining Risk in Data-Poor Fisheries* to estimate the effective number of attributes used for each variable. - Classify each variable as additive or multiplicative in nature. When an additive model is used for a variable, it is assumed that each attribute contributes to a fraction of risk proportional to its weight (see following instruction on weighting). Therefore, adding all attribute contributions to risk gives the full risk associated with the variable. When a multiplicative model is used for a variable, it is assumed that each attribute’s contribution is affected by the contributions of other attributes. Simply, if risk from one or more attributes is absent, overall risk associated with the variable would be absent as well even when high risk is associated with other attributes. This model is often used when attributes measured operate in a sequence or are probabilistic (e.g. Likelihood). By default, response variables are additive and stressor variables are multiplicative in the templates. Different models can be used for each stressor, though care should be taken to interpret results appropriately given the added complexity of the analysis. - Set criteria for low, medium, and high risk for each variable. This consists of two percentile cutoffs, below the first is low, above the second is high, and between them is medium risk. These cut-offs can be chosen as any percentile provided they are symmetric (i.e. low and high categories are of equal range). The template defaults assume equally sized categories. - Assign weights to each attribute. Weights can take any numerical value and are only important relative to weights of other attributes for a single species or endpoint. Weights can differ within an attribute but across species. The template defaults to equal weighting. - Score attributes for all species. Attribute scores should take values between 1 and 3. These values should fall within intended categories based on the percentile cut-offs assigned above. Defaults correspond to 1 = low risk, 2 = medium risk, 3 = high risk.

### 2. Template completion

- Download one alpha template and one beta csv template. - Open the alpha template. The top four rows have six sets of values to input. a) Additive or Multiplicative attributes? Acceptable inputs: Additive, Multiplicative b) Low and high percentile cut-offs for attribute scoring. Acceptable inputs: any two fractions that as cut-offs produce a symmetric distribution of score ranges. Numerators and denominators must include a decimal. Values must remain in fraction form (i.e. do not give the decimal equivalent of the fraction). c) Number of attributes. Acceptable input: a whole number corresponding to the number of attribute columns. d) Low and high thresholds. Acceptable inputs: The second value must be larger than the first. Numerators and denominators must include a decimal. Values must remain in fraction form (i.e. do not give the decimal equivalent of the fraction). These thresholds determine the risk categories following Vulnerability scoring. e) Scoring in reversed risk order? Acceptable inputs: Y, N. In some cases (e.g. Productivity) the attributes of the alpha variable may be scored such that high values represent low risk. If scores input in the template were scored in this way, assign Y to this field. If high values correspond to high risk, assign N to this field. f) Axis label. Acceptable inputs: any x-axis label for the resulting plot, preferably the name of the alpha variable. - Row 5 must be left empty. Row 6 is the dataset header, and values in these cells can be changed without affecting the analysis. - Columns must be organized accordingly: column 1 is for higher level organization of species and will not be output in results. Column 2 is the list of species and will be reported in results alongside Vulnerability scores and risk categories. Input your list of species in column 2. No input is required for column 1 unless it is helpful for your organization. Blank rows can be included in between species or chunks of species for aesthetics without affecting the analysis provided the blank rows are placed consistently for all templates so that species fall on the same row. - Columns 3+ are for attribute scores and weights. Add or remove columns to rows 6+ to add or subtract attributes. Two empty columns must be kept between attribute scores and attribute weights. Attribute weight columns must be in the same order, left to right, as the attribute columns. For example, for 5 attributes, columns 1 and 2 would report group (optional) and species (or endpoint generally). Columns 3-7 would report attribute scores. Columns 8-9 would be blank. Columns 10-14 would report attribute weights. When an attribute is unscored due to lack of data or irrelevance for the species, both the attribute score and weight cells should be left blank. Default attribute scores are randomly chosen between 1 and 3. Default weights are equal. - Save the alpha template as alpha xxx.csv where xxx is any string you choose. The file must be saved in UTF-8 format. Note: Microsoft Office for Mac incorrectly encodes the UTF-8 format, so upload errors may be a result of incorrect encoding. Use LibreOffice, Google Sheets, or Numbers on a Mac. Microsoft Office works correctly on a PC for csv encoding. - Open the beta template. Like the alpha template, the top four rows have six sets of values to input. These sets of values can be entered according to the guidelines for the alpha template above except for (e). Here the entry differs: Compound model? Acceptable inputs: Y, N. This entry refers to whether the stressor is statistically standardized by the expected mean or the expected minimum. By default, this value should be N for the first beta template to yield a comparable analysis to the rPSA for a single stressor. Subsequent stressors can be treated as compounding (increasing risk with more stressors – Y) or not (N) by completing additional beta templates for each stressor. - The same rules apply for column and row formatting and data entry for both beta and alpha templates. It is recommended to rank stressors in order of impact, with the highest impact stressor entered in the first beta template, and the lowest impact stressor entered in the last beta template. All values can differ between stressors except for (d) low and high thresholds. These thresholds will be the same across all templates, including the alpha template, as the thresholds are applied to Vulnerability scores at the end of the analysis. The software is setup to take the threshold values from the alpha template, so modifying the thresholds in the beta templates will not change results. - Save each beta template as beta1 xxx.csv, beta2 xxx.csv, beta3 xxx.csv, etc. in the same folder as you saved the completed alpha template. Ordering of these completed templates matters in the upload stage, as all files are selected simultaneously. Therefore, in the folder, files must appear in the following order: alpha xxx.csv, beta1 xxx.csv, beta2 xxx.csv, beta3 xxx.csv, etc.

### 3 EcoRAMS analysis

- Navigate to the main page of EcoRAMS.net. - Click on the ‘Choose Files’ button after which a file browser window will appear. Navigate to the folder hosting your completed templates. The order the files appear is the order they will be uploaded and should be in the order described above. Use ctrl (or cmd) + select or shift select to select all files to be analyzed. - After opening these files, the homepage will read the number of files selected. Click ‘Submit’ to analyze your data. - After a few seconds, a display page will appear with results. Results will appear in order of the stressors uploaded with stressor 1 corresponding to beta1 xxx.csv. These are single stressor results for each stressor standardized by the expected mean. Each plot will precede a list of all species and their associated probabilistic Vulnerability score between 0 (lowest) and 1 (highest) and their associated risk category determined by the thresholds set. The final result is the multiple stressor result. - All figures can be downloaded by saving the image with a right click, and the data tables can be copied and pasted directly into any format like a csv file.

## References

Battista, W., Karr, K., Sarto, N. & Fujita, R. (2017). Comprehensive assessment of risk to ecosystems (care): A cumulative ecosystem risk assessment tool. Fisheries Research, 185, 115–129.

Council, N. R. et al. (2009). Science and decisions: Advancing risk assessment.

FAO. (2020). The state of world fisheries and aquaculture (sofia): 2020.

Finkbeiner, E. M., Bennett, N. J., Frawley, T. H., Mason, J. G., Briscoe, D. K., Brooks, C. M., Ng, C. A., Ourens, R., Seto, K., Switzer Swanson, S. et al. (2017). Reconstructing overfishing: Moving beyond malthus for effective and equitable solutions. Fish and Fisheries, 18 (6), 1180–1191.

Grewelle, R. E., Mansfield, E., Micheli, F. & De Leo, G. (2021). Redefining risk in data-poor fisheries. Fish and Fisheries.

Halpern, B. S., Kappel, C. V., Selkoe, K. A., Micheli, F., Ebert, C. M., Kontgis, C., Crain, C. M., Martone, R. G., Shearer, C. & Teck, S. J. (2009). Mapping cumulative human impacts to california current marine ecosystems. Conservation letters, 2 (3), 138–148.

Hazen, L., Le Cornu, E., Zerbe, A., Martone, R., Erickson, A. L. & Crowder, L. B. (2016). Translating sustainable seafood frameworks to assess the implementation of ecosystem-based fisheries management. Fisheries Research, 182, 149–157.

Hobday, A., Smith, A., Stobutzki, I., Bulman, C., Daley, R., Dambacher, J., Deng, R., Dowdney, J., Fuller, M., Furlani, D. et al. (2011). Ecological risk assessment for the effects of fishing. Fisheries Research, 108 (2-3), 372–384.

Hobday, A., Smith, A., Webb, H., Daley, R., Wayte, S., Bulman, C., Dowdney, J., Williams, A., Sporcic, M., Dambacher, J. et al. (2007). Ecological risk assessment for the effects of fishing: Methodology. report r04/1072 for the australian fisheries management authority.

Hope, B. K. (2006). An examination of ecological risk assessment and management practices. Environment International, 32 (8), 983–995.

Koller, G. (2005). Risk assessment and decision making in business and industry: A practical guide. CRC Press.

Marine Stewardship Council. (2019). The msc annual report 2018-2019.

Micheli, F., De Leo, G., Butner, C., Martone, R. G. & Shester, G. (2014). A risk-based framework for assessing the cumulative impact of multiple fisheries. Biological Conservation, 176, 224–235.

Mora, C., Tittensor, D. P., Adl, S., Simpson, A. G. & Worm, B. (2011). How many species are there on earth and in the ocean? PLoS biology, 9 (8), e1001127.

Norton, S. B., Rodier, D. J., van der Schalie, W. H., Wood, W. P., Slimak, M. W. & Gentile, J. H. (1992). A framework for ecological risk assessment at the epa. Environmental toxicology and chemistry, 11 (12), 1663–1672.

Oestreich, W. K., Frawley, T. H., Mansfield, E. J., Green, K. M., Green, S. J., Naggea, J., Selgrath, J. C., Swanson, S. S., Urteaga, J., White, T. D. et al. (2019). The impact of environmental change on small-scale fishing communities: Moving beyond adaptive capacity to community response. Predicting future oceans (pp. 271–282). Elsevier.

Pilling, G. M., Apostolaki, P., Failler, P., Floros, C., Large, P. A., Morales-Nin, B., Reglero, P., Stergiou, K. I. & Tsikliras, A. C. (2009). Assessment and management of data-poor fisheries. Advances in fisheries science, 50, 280–305.

Pontón-Cevallos, J. F., Bruneel, S., Marin Jarrin, J. R., Ramirez-González, J., Bermúdez-Monsalve, J. R. & Goethals, P. L. (2020). Vulnerability and decision-making in multispecies fisheries: A risk assessment of bacalao (mycteroperca olfax) and related species in the galapagos’ handline fishery. Sustainability, 12 (17), 6931.

Ravindra, K. & Mor, S. (2019). Distribution and health risk assessment of arsenic and selected heavy metals in groundwater of chandigarh, india. Environmental pollution, 250, 820–830.

Samhouri, J. F., Ramanujam, E., Bizzarro, J. J., Carter, H., Sayce, K. & Shen, S. (2019). An ecosystem-based risk assessment for california fisheries co-developed by scientists, managers, and stakeholders. Biological Conservation, 231, 103–121.

Stobutzki, I., Miller, M. & Brewer, D. (2001). Sustainability of fishery bycatch: A process for assessing highly diverse and numerous bycatch. Environmental Conservation, 28 (2), 167–181.

Tannenbaum, L. V., Johnson, M. S. & Bazar, M. (2003). Application of the hazard quotient method in remedial decisions: A comparison of human and ecological risk assessments. Human and Ecological Risk Assessment, 9 (1), 387–401.

Townsend, H., Harvey, C. J., deReynier, Y., Davis, D., Zador, S., Gaichas, S., Weijerman, M., Hazen, E. L. & Kaplan, I. C. (2019). Progress on implementing ecosystem-based fisheries management in the us through the use of ecosystem models and analysis. Frontiers in Marine Science, 6, 641.

Van den Brink, P. J., Choung, C. B., Landis, W., Mayer-Pinto, M., Pettigrove, V., Scanes, P., Smith, R. & Stauber, J. (2016). New approaches to the ecological risk assessment of multiple stressors. Marine and Freshwater Research, 67 (4), 429–439.

World Health Organization et al. (2020). Risk assessment.

Zio, E. (2018). The future of risk assessment. Reliability Engineering & System Safety, 177, 176–190.

